# Separation of coiled-coil structures in lamin A/C is required for elongation of the filament

**DOI:** 10.1101/2020.09.29.317933

**Authors:** Jinsook Ahn, Soyeon Jeong, So-mi Kang, Inseong Jo, Bum-Joon Park, Nam-Chul Ha

## Abstract

Intermediate filaments (IFs) commonly have structural elements of a central α-helical coiled-coil domain consisting of coil 1a, coil 1b, coil 2, and their flanking linkers. Recently, crystal structure of a long lamin A/C fragment was determined and showed detailed features of a tetrameric unit. The structure further suggested a new binding mode between tetramers, designated eA22, where a parallel overlap of coil 1a and coil 2 is the key interaction. In this study, we investigated the biochemical effects of genetic mutations causing human diseases, focusing on the eA22 interaction. The mutant proteins exhibited either weakened or augmented interactions between coil 1a and coil 2. The ensuing biochemical results indicated that the interaction requires the separation of the coiled-coils in N-terminal of coil 1a and C-terminal of coil 2, coupled with the structural transition in the central α-helical rod domain. This study provides insight into the role of coil 1a as a molecular regulator in elongation of the IF proteins.

## INTRODUCTION

Intermediate filament (IF) proteins provide vital mechanical support in higher eukaryotic cells, with various physical properties when they form filamentous polymers (Holaska, Wilson et al., 2002, Lopez-Soler, Moir et al., 2001, Stuurman, Heins et al., 1998). The monomeric units of all IF proteins share a tripartite structural organization: a central α-helical rod domain flanked by non-α helical N-terminal head and C-terminal tail regions (Gu, Troncoso et al., 2004, Herrmann, Bar et al., 2007, Herrmann, Haner et al., 1999, Parry & Steinert, 1999). The α-helical rod domain mostly consists of multiple heptads and hendecads, imparting a high propensity to form parallel coiled-coil dimers with different super-helical pitches. The α-helical rod domain is divided into three segments (coil 1a, coil 1b, and coil 2) by flanking linkers (L1 and L12, respectively) lacking periodic repetitions (Parry & Steinert, 1999). The stutter region within coil 2 shows a different periodic rule from the adjacent areas (Brown, Cohen et al., 1996, Lupas, 1996). Two IF consensus sequence motifs are found at both end regions of the central rod domain (Fig. 1A), which are highly conserved among IF proteins and play a crucial role in forming the filamentous structure (Burkhard, Stetefeld et al., 2001, Herrmann & Strelkov, 2011, Herrmann, Strelkov et al., 2000, Kouklis, Traub et al., 1992, Schaffeld, Herrmann et al., 2001). The α-helical rod domains are assembled to produce a long filamentous structure with 10-nm thickness in cytoplasmic IF proteins (Goldman, Gruenbaum et al., 2002, Hess, Voss et al., 2002, Turgay, Eibauer et al., 2017).

**Figure 1.**
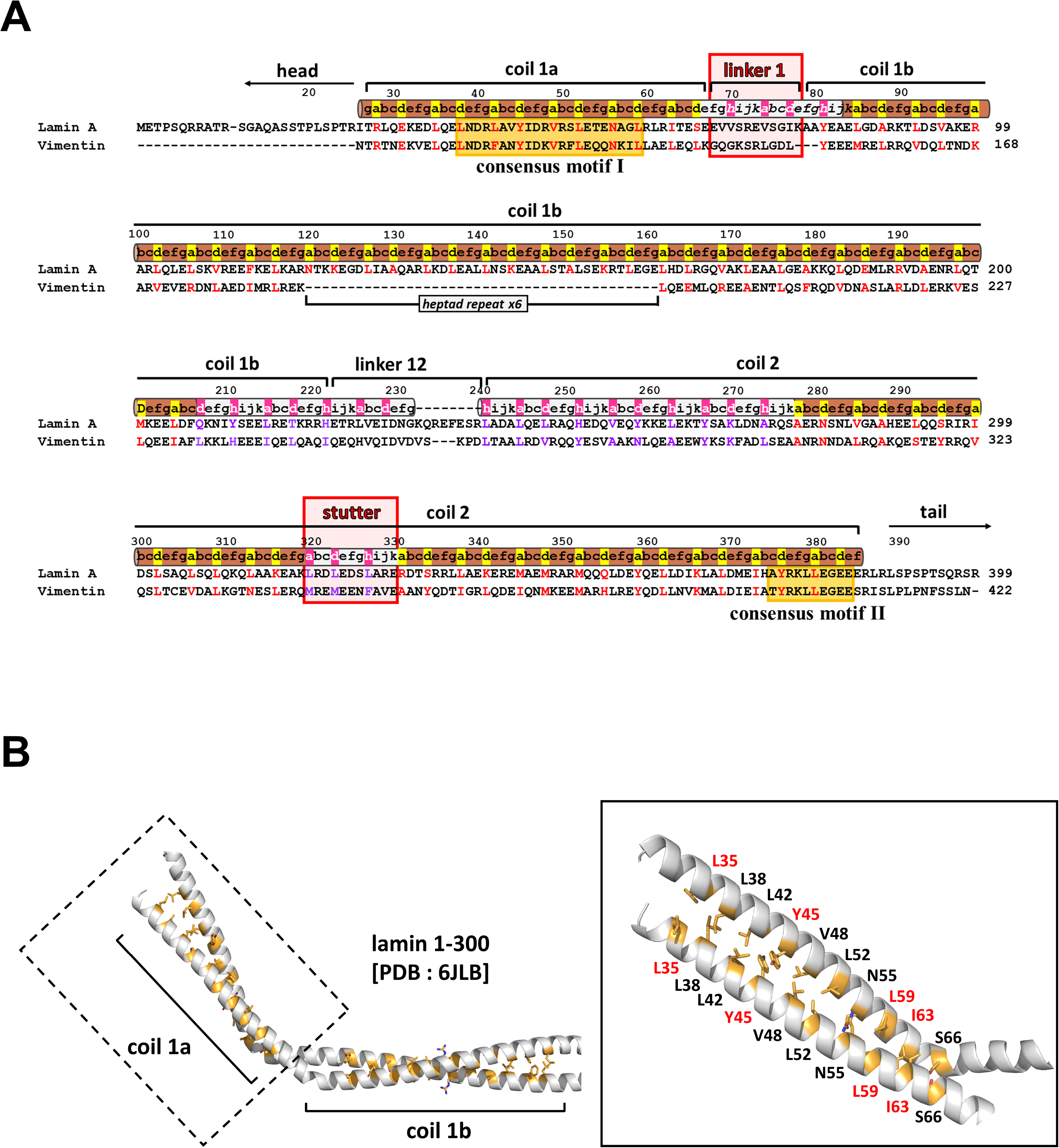
Kinked coiled-coil structure of the N-terminal region of coil 1a and coil 1b parts in lamin A/C (PDB 6JLB (Ahn et al., 2019)) (A) Sequence alignment of the central rod domain in lamin A/C and vimentin proteins. Pairwise sequence alignment of lamin (human lamin A/C; P02545.1) and putative IF proteins (human vimentin; NP_003371.2). Sequence periodicity in the heptad (brown) or hendecad (grey) repeat, which was previously predicted (Chernyatina, Guzenko et al., 2015, Parry, 2006), and revealed by the crystal structure of lamin 1-300 (Ahn et al., 2019), is displayed above the sequences. The yellow boxes indicate highly conserved residues (consensus motif I and II). Linker 1 and stutter regions are highlighted in red boxes. The *a*, *d*, and *h* residues are highlighted in red for heptad and purple for hendecad repeats. (B) The ribbon diagram presents an N-terminal area composed of coil 1a, linker 1, and half of coil 1b. The coil 1a region is enlarged in the box (*right*). The inter-helical hydrophobic residues at the a and d positions of the heptad repeat are shown as yellow sticks, and the residues for further mutational studies are labelled in red.

Nuclear IF lamin A/C is essential in formation and maintenance of the nuclear structure by providing rigidity and flexibility underneath the inner nuclear membranes (Burke & Stewart, 2013, Chojnowski, Ong et al., 2015, Koster, Weitz et al., 2015). The parallel coiled-coil dimers of lamin A/C, as a fundamental building block, assemble laterally and longitudinally into high-order filamentous structures (Koster et al., 2015, Parry, Strelkov et al., 2007). Recent *in situ* cryo-electron tomography (cryo-ET) images visualized the 3.5-nm-thick filament of lamin A/C, decorated with an Ig-like fold domain at both sides with 20-nm longitudinal intervals, unlike 10-nm-thick cytosolic IF proteins (Turgay et al., 2017). The assembly model of lamin A/C was substantially detailed at higher resolution by the crystal structure of the N-terminal half fragment of lamin A/C, called the lamin 300 fragment. The crystal structure presented the structural features of the so-called ‘A11 tetramer,’ which consists of two coiled-coil dimers in an anti-parallel manner by overlapping the coil 1b regions (Ahn, Jo et al., 2019). The crystal structure and biochemical analyses indicate ‘eA22 binding mode’ for joining the A11 tetramers. The parallel overlapping between the C-terminal region of coil 2 and the coil 1a in the A11 tetramer was crucial for the eA22 interaction to form the eA22 binding mode. However, little is understood of the molecular mechanism of eA22 interaction for elongation of IF proteins.

Many genetic mutations in the lamin A/C gene result in human diseases, such as Emery Dreifuss muscular dystrophy (EDMD), dilated cardiomyopathy (DCM), Dunnigan-type familial partial lipodystrophy (FPLD), and Hutchinson-Gilford progeria syndrome (HPGS) (Butin-Israeli, Adam et al., 2012, Dittmer & Misteli, 2011, Liu & Zhou, 2008, Muchir, Medioni et al., 2004). The genetic variations causing diseases were highly associated with two IF consensus motifs interfacing parallel overlapping region between coil 1a and coil 2 (Ahn et al., 2019, Burkhard et al., 2001, Herrmann & Strelkov, 2011, Herrmann et al., 2000, Kouklis et al., 1992, Schaffeld et al., 2001). In this study, we first investigate the role of eA22 interaction in lamin-related diseases, focusing on EDMD and DCM. Then, we propose a binding model for the eA22 interaction remodelling the coiled-coil interactions, coupled with the structural transition in the central α-helical rod domain.

## RESULTS

### Mutations of EDMD and DCM altered the eA22 interaction

In a previous study, we investigated the Y45C and L59R mutations in coil 1a in terms of eA22 interaction (Ahn et al., 2019). Mutation of Y45C in EDMD abolished the eA22 interaction, while the L59R mutation in DCM increased the eA22 interaction. These observations suggest that EDMD and DCM are associated with increased and decreased eA22 interaction, respectively, compared to wild type lamin A/C. To test the hypothesis for generalization to other mutations in EDMD and DCM, we investigated two additional mutations, L35V in DCM and I63S in EDMD. All four mutations (L35V, Y45C, L59R, and I63S) were in the *a* or *d* position of coil 1a in the crystal structure (Ahn et al., 2019), which potentially affects the coiled-coil interaction in the coiled-coil dimeric unit (Fig. 1) (Dittmer & Misteli, 2011, Kang, Yoon et al., 2018, Maraldi, Squarzoni et al., 2005, Muchir et al., 2004, Strelkov, Schumacher et al., 2004).

We purified four mutant proteins based on the lamin 300 fragment (residues 1-300) as the coil 1a-containing proteins to examine the eA22 interaction. The mutant proteins were expressed as a dimeric unit in solution, like the wild type fragment. We performed the His-tag pull-down assay to evaluate the strength of the eA22 interaction, as previously described (Ahn et al., 2019). The His-tagged mutant lamin 300 fragments were incubated with the C-terminal part of coil 2 (residues 250-400) in a low-salt buffer containing 50 mM NaCl or a high-salt buffer containing 150 mM NaCl. The L35V mutation in DCM and the Y45C mutation in EDMD reduced the binding strength of eA22 interaction in both buffers (Fig. 2). In contrast, L59R in DCM and I63S in EDMD increased the eA22 interactions in both buffer conditions (Fig. 2). Our findings confirmed that altered eA22 interaction is involved in EDMD and DCM. However, the results suggest that EDMD and DCM cannot be distinguished only by the higher or lower strength of the eA22 interaction.

**Figure 2.**
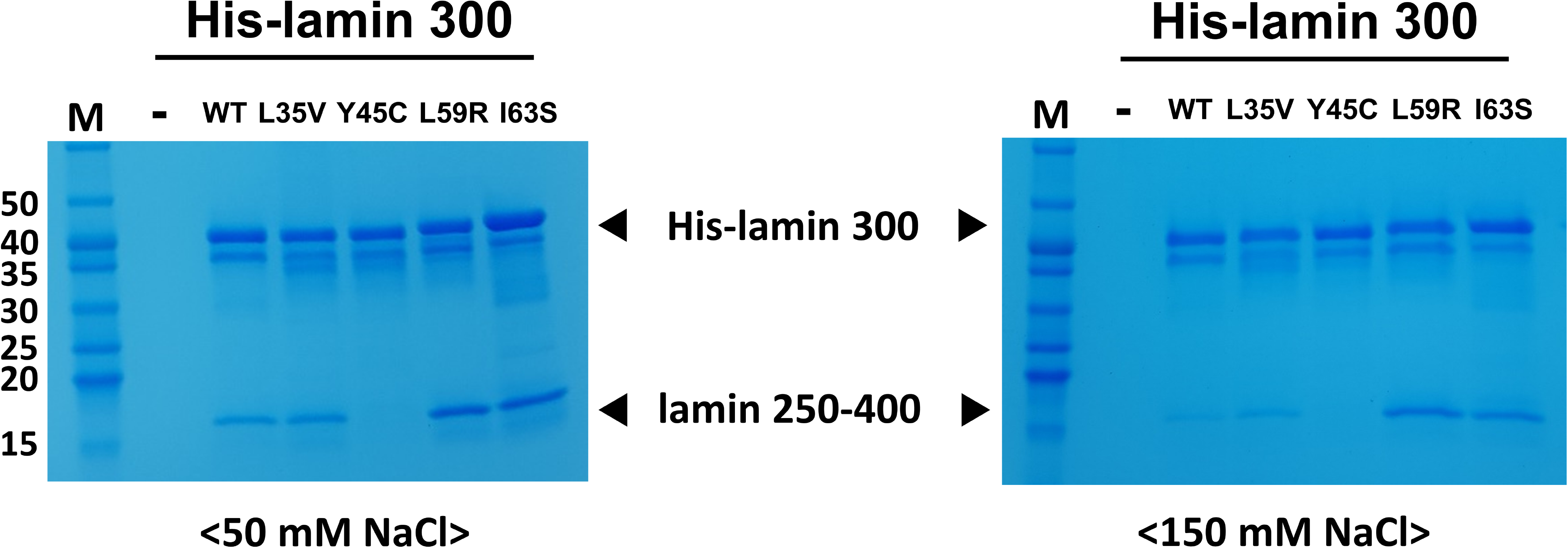
The strength of the eA22 interaction in the four mutations related to laminopathies. The binding affinity between the lamin 250-400 (C-terminal of coil 2) and His-lamin 1-300 variants (WT, L35V, Y45C, L59R, and I63S) was analysed by the His pull-down assays. Tag-free lamin 250-400 fragments were incubated on empty (-) or His-tagged lamin-bound (WT, L35V, Y45C, L59R, and I63S) Ni-NTA resins. The resins were pre-equilibrated and washed with 20 mM Tris-HCl (pH 8.0) buffer containing 50 mM (*left*) or 150 mM NaCl (*right*). After washing, the bound proteins were analysed by SDS-PAGE. Molecular weights (kDa) of the marker (M) are labeled on the left.

Next, we employed isothermal titration calorimetry (ITC) to analyse the binding affinities quantitatively. Similar K_D_ values for wild type and L59R mutant were measured and compared with values previously reported (Ahn et al., 2019). We included K_D_ values for the rest of the mutant proteins in this study. Based on the measured K_D_ values, the L59R and I63S mutants increased the eA22 interaction by ~56 fold and ~2 fold, respectively (Fig. 3). In contrast, the L35V and Y45C mutations decreased the eA22 interaction by ~2 fold and ~5 fold, respectively. These results are in good agreement with the results from the pull-down assays shown in Fig. 2.

**Figure 3.**
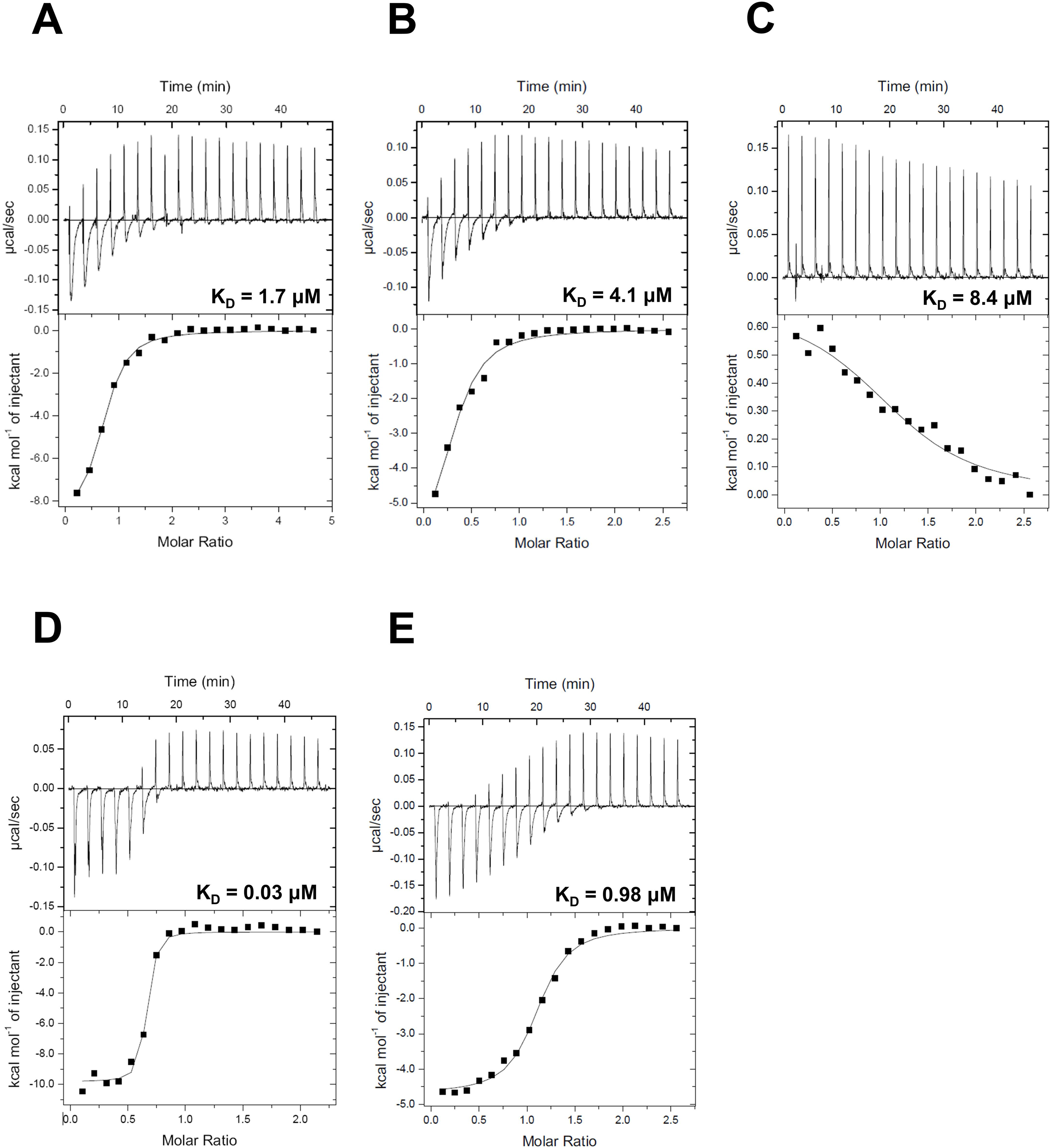
Isothermal titration calorimetry analysis for the binding affinity between the lamin 300 fragments (A, WT; B, L35V; C, Y45C; D, L59R; E, I63S) and the coil 2 fragment (residues 250-400) The top panels represent the raw measured heat changes (μcal/s) as a series of peaks corresponding to a function of time resulting from the titration of each lamin 300 fragment (20 μM; 370 μL) with 19 injections of the coil 2 fragment (160 μM; 2 μL per one injection). The bottom panels show the titration isotherm resulting from the raw data in the top panels.

To examine the effects of the mutations on nuclear shape and distribution of lamin A/C in cells, we overexpressed the mutant lamin genes in the HEK293 cell lines. Lamin A L35V and Y45C, with a lower eA22 interaction, diffused to the cytosol, forming localized aggregations, indicating a lack of lamin filament formation at the nuclear membrane. However, lamin A L59R and I63S, with higher eA22 interactions, presented strong lamin aggregates at the peripheral region of the nuclear envelopes, indicating tangled filament formation (Fig. 4). The results from Y45C and L59R were similar to previous observations (Ahn et al., 2019), and all the mutations produced the same results in the human bone osteosarcoma epithelial (U2OS) cell lines (Fig EV1). Our findings confirm the previous proposition that both stronger and weaker eA22 interactions result in adverse effects on the formation of robust nuclear structures than the proper eA22 interaction of wild type lamin.

**Figure 4.**
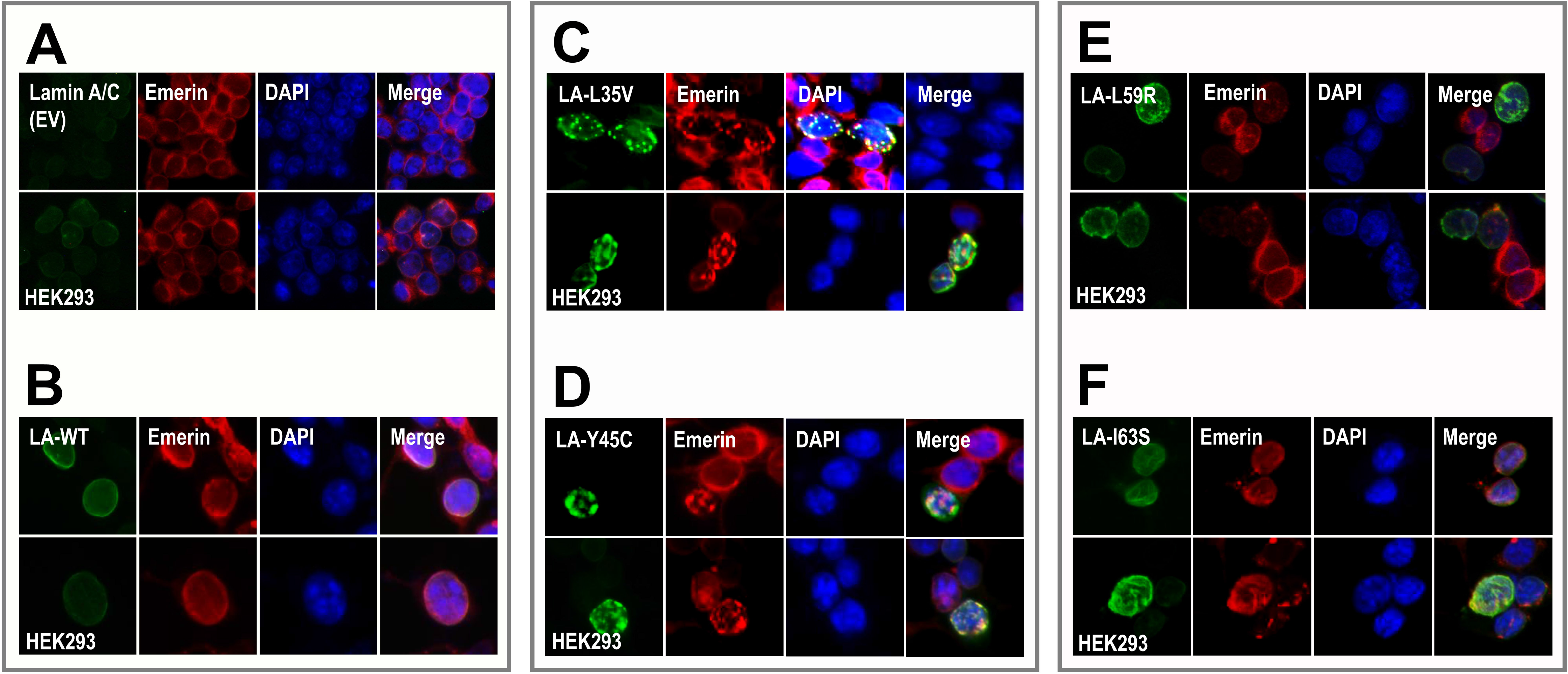
Nuclear shapes and distribution of the wild type and mutants (L35V, Y45C, L59R, and I63S) of lamin A/C. The immunofluorescence assay visualized nuclear morphology after the transfection of wild type or variants (L35V, Y45C, L59R, and I63S) of lamin A/C into HEK293 cells. For visualization of the nuclear membrane, cells were stained with the corresponding antibodies (lamin A/C; green, emerin; red) and DAPI for DNA (blue). The merged images of lamin A/C, emerin, and DNA are displayed on the left (merge).

### The eA22 interaction requires separation of the coiled-coil dimer in the coil 1a region

To explain the mutations affecting the eA22 interaction from a structural view, we closely examine the mutations in the dimeric structure of the lamin 300 fragment (Fig. EV2) (Ahn et al., 2019). The modelled structure of the L35V mutant suggests that the mutation would stabilize the coiled-coil dimer at the coil 1a region because the distance between coiled-coils would decrease. According to the ΔΔGu (Ala) values indicating the stability of the coiled-coil structure at the *a* and *d* positions (Meier, Padilla et al., 2009, Tripet, Wagschal et al., 2000), valine stabilizes the coiled-coil structure of coil 1a compared to the leucine residue (Fig. EV2A).

Tyr45 is conserved among vimentin as well as lamin A and B families. Tyrosine residues at the *a* and *d* positions destabilize the coiled-coil structure in general (Ahn et al., 2019, Tripet et al., 2000). Thus, the Y45C mutation was suggested to stabilize the coiled-coil interaction of coil 1a because the cysteine residue is a better fit at the *d* position for coiled-coil formation (Fig. EV2B) (Ahn et al., 2019). Conversely, loss of their hydrophobicity in the *a* or *d* heptad position seems to be the reason why the mutations of L59R and I63S weakened the coiled-coil dimer structure. In particular, the modelled structure of the L59R mutant showed that Arg59 sterically clashed with and repulsed each other due to their bulky and positive charge (Fig. EV2C) (Ahn et al., 2019).

We explored the structural features of mutant lamin proteins focusing on the coil 1a region using circular dichroism (CD) spectroscopy. The CD spectra demonstrate the propensity for the secondary structure of the proteins. Since the separated α-helices in solution would not be as stable as the coiled-coil dimers, the α-helical tendency of the coil 1a region is a good indicator of the stability of the coiled-coil dimers (Meier et al., 2009). To exclude the background α-helicity in the resting region, we used shorter fragments covering residues 1-125 containing the N-terminal head, coil 1a, L1, and a small portion of coil 1b instead of the lamin 300 fragment. The shorter fragments harbouring the mutations exhibited similar results to those with the longer lamin 300 fragments in terms of alteration of eA22 interaction (Fig. 5 and Fig. EV3). The CD spectra of the shorter fragments showed that the Y45C mutant protein has the highest α-helical content of 54%, indicating the largest increase in stability of coiled-coil dimers (Fig. 5B; *red*). The L59R mutant has the lowest α-helical content of 16%, presumably indicating substantial disruption of coiled-coil dimers (Fig. 5B; *green*). The L35V and I63S proteins showed slight differences in terms of α-helical content, which agree well with the relatively mild affinity changes to the coil 2 region (Fig. 5B; *yellow* and *purple*). Thus, our findings showed that the degree of α-helix tendency is correlated to the results for eA22 interactions: smaller α-helical content or weaker coiled-coil interaction produce stronger eA22 interactions.

**Figure 5.**
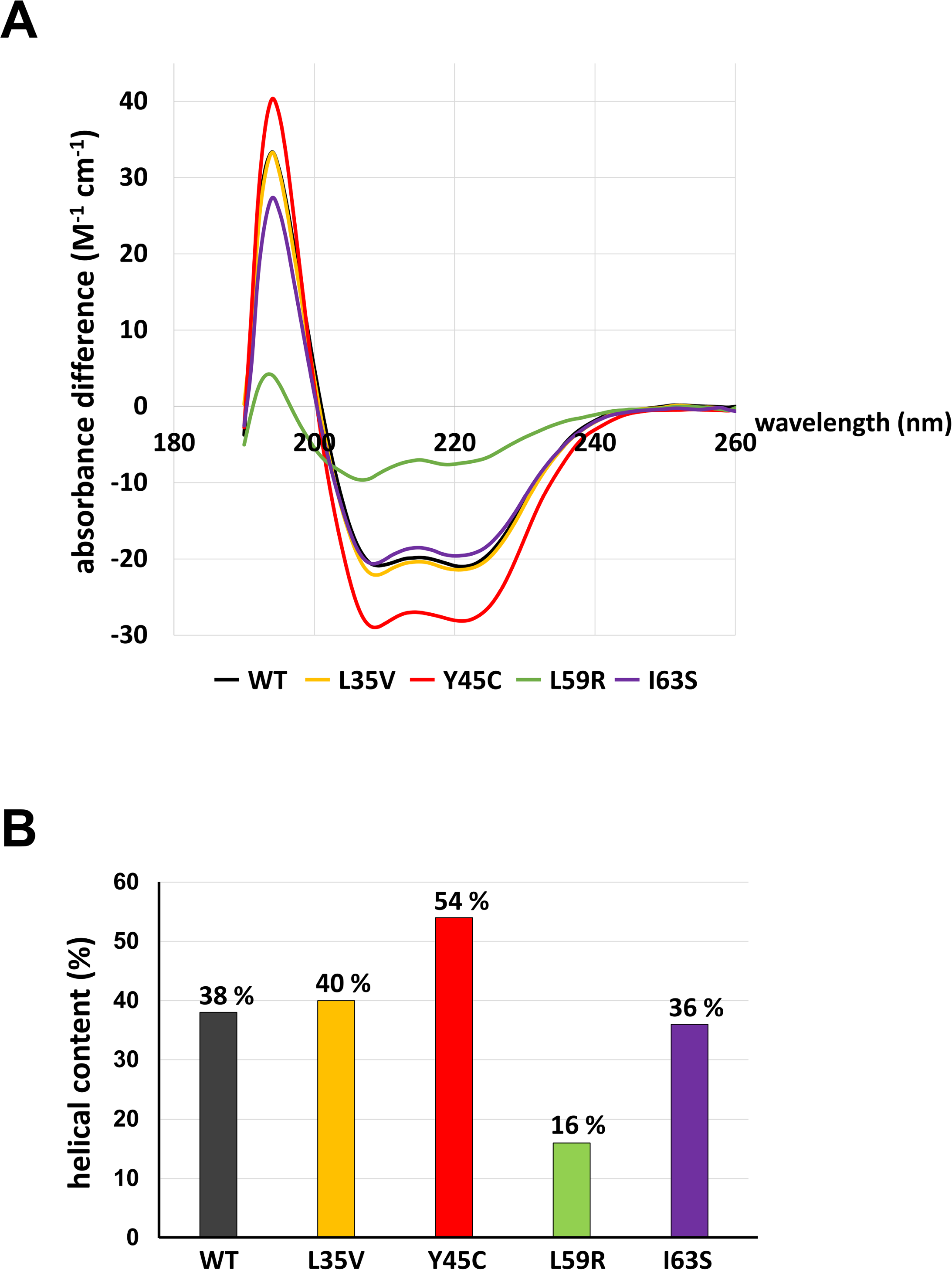
The α-helical contents of lamin 1-125 fragments (WT, L35V, Y45C, L59R, and I63S) (A) The CD spectra were recorded at a protein concentration of 1 mg/mL. Wild type and variant proteins are coloured differently (WT; grey, L35V; yellow, Y45C, L59R; green, I63S; violet). Estimated helical contents of lamin 1-125 fragments are presented in (B).

### Leu38 and Leu42 at the N-terminal end region of coil 1a are vital in interaction with the coil 2 region

The L38C and L42C mutant proteins in the shorter N-terminal fragment (residues 1-125) were generated to analyse the effect of the complete prevention of separation of the coiled-coils in coil 1a on the formation of the eA22 interaction. Since Leu38 and Leu42 residues are at the *a* and *d* positions and in close contact in the coiled-coil structure, changing these residues to cysteine would form a disulfide bond under oxidized conditions, resulting in the prevention of separation of the coil-coil structure (Fig. 1). Unexpectedly, mutant proteins did not bind to the coil 2 fragment under either reduced or oxidized conditions (Fig. EV4). These findings indicate that Leu38 and Leu42 are essential for the eA22 interaction, presumably to bind with the coil 2 region.

### The eA22 interaction requires separation of coiled-coil dimers in the C-terminal region of coil 2

Using the same strategy as in the coil 1a part, we explored the role of the coil 2 part in the eA22 interaction. The C-terminal region of coil 2 can be divided into two subparts by the stutter region (residues 320-330; Fig. 1A). The stutter region is essential for elongation of the lamin and the other cytosolic IF proteins (Parry & Steinert, 1999, Smith, Strelkov et al., 2002). The crystal structures of these regions of lamin B1 and vimentin represented that the C-terminal region of the coil 2 containing the stutter region maintained the coiled-coil α-helical conformation with the corresponding coiled-coil patterns (Brown et al., 1996, Lupas, 1996, Nicolet, Herrmann et al., 2010).

To test if the separation of the coiled-coils is required during the eA22 interaction, we introduced four single mutations at the *a* or *d* positions of the heptad or hendecad pattern in the coil 2 fragment. Each residue was replaced with cysteine to form a disulfide bond in the oxidized condition, resulting in blocked separation of the coiled-coil dimer. Two residues were selected before the stutter region, and the others after the stutter region. Of four mutants (Y267C, K316C, L362C, and L380C), the L380C mutation abolished the eA22 interaction with the coil 1a fragment even in the reduced condition, indicating that Leu380 is essential in the interaction with the coil 1a part (Fig. 6A and Fig. EV5A). The subsequent L380A mutant protein confirmed the importance of Leu380 in the eA22 interaction (Fig. 6B and Fig. EV5B).

**Figure 6.**
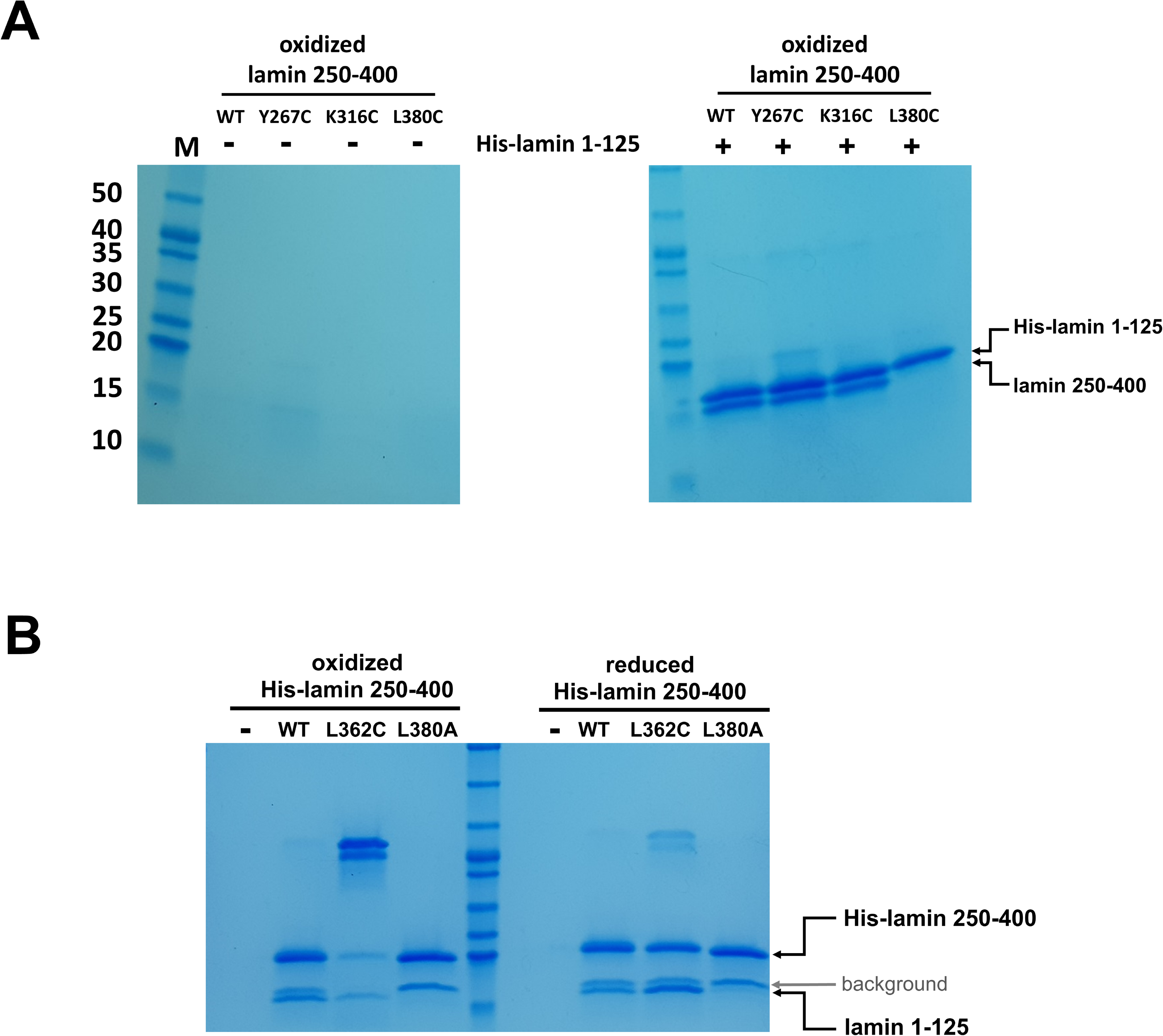
Altered eA22 interaction depending on CC dimer separation of the coil 2 fragment. (A) The wild type of His-tagged lamin 1-125 was immobilized on Ni-NTA resin. Lamin 250-400 fragments (WT, Y267C, K316C, and L380C) were incubated on empty (-) or His-lamin-bound (+) Ni-NTA resins under the non-reducing condition. The bound proteins were analysed using SDS-PAGE. (B) The His-tagged lamin 250-400 fragments (WT, L362C, and L380A) were immobilized on Ni-NTA resin with the lamin 1-125 fragment. The mixed samples were incubated under the reducing or non-reducing condition. A degraded His-lamin 250-400 fragment band is shown between the molecular sizes of 15 and 20 kDa in SDS-PAGE (background, grey arrow). The used proteins for binding affinity are shown in Fig. EV5.

Importantly, the L362C mutation, located after the stutter region, abolished the eA22 interaction only under the oxidized condition allowing the disulfide bond that prevents rearrangement of the coiled-coil structure of the C-terminal region of coil 2 (Fig. 6B and Fig. EV5B). In contrast, the other two mutations (Y267C and K316C), located before the stutter region, did not affect the eA22 interaction under either reduced or oxidized conditions (Fig. 6A and Fig. EV5A). These findings indicate that separation of the coiled-coils at the C-terminal region of coil 2 after the stutter region is required for the eA22 interaction.

## DISCUSSION

To account for the conformational transition of coiled-coil dimers both in the N-terminal of coil 1a and C-terminal of coil 2 parts in the eA22 interaction, we closely examined the structures of the lamin 300 fragments and vimentin (residues 99 – 189), both of which contain coil 1a, coil 1b, and flanking linker L1 (Fig. 1B and Fig. EV6) (Ahn et al., 2019, Meier et al., 2009). Both structures exhibited all α-helical conformation in the coil 1a and coil 1b regions, including the flanking linker that did not follow the heptad rule. However, the two structures presented quite a different arrangement in terms of coiled-coil dimer formation and bending of α-helices. We noted that the abrupt bending was found at the linker L1 region connecting coil 1a to coil 1b in the continuous coiled-coil structure, and that linker L1 does not follow the heptad rule (Fig. 1B). In contrast, no abrupt kink was found in the vimentin structure with separation of the coiled-coil in the coil 1a region, also caused by the broken heptad rule at the linker (Fig. EV6). The structural comparison suggests that the kink at the linker compensates for the broken heptad rule, resulting in the formation of stable coiled-coil interaction throughout coil 1a and coil 1b regions. Otherwise, the coiled-coil would be separated without the kink at the linker region. Thus, our findings suggest a dynamic equilibrium between two conformations: kinked and coiled-coil (CC) separated conformation

Next, we applied a similar mechanism in the C-terminal region of coil 2. We showed that the eA22 interaction also requires the CC-separation of the C-terminal part of coil 2. We observed that blocking of the coiled-coil separation after the stutter region abolished the eA22 interaction (Fig. 6). Significant conformational rearrangement, accompanied by coiled-coil separation, would occur at the C-terminal part of the coil 2 for the eA22 interaction. The stutter region does not follow the heptad rule, as in linker L1, and the rest of the coil 2 region loosely follows the heptad rule. We further expect that the four-helix bundle model, previously proposed by Herrmann and Strelkov et al. (Herrmann & Strelkov, 2011, Kapinos, Burkhard et al., 2011, Kapinos, Schumacher et al., 2010), represent the resulting complex structure of coil 1a and coil 2.

Based on the dynamic conformational equilibrium, we propose an assembly model for the eA22 interaction in the elongation process of IFs (Fig. 7). Until the cognate partner proteins appear, the A11 tetramer which formed transiently in lamin A/C has the dynamic equilibrium between two conformations; kinked and coiled-coil (CC) separated conformation. Then, two A11 tetramers produce the eA22 interaction by the straight and separated conformation for elongation of the filament. This assembly model has many advantages in accounting for selective assembly of the IF members to identify the appropriate binding partners among more than 67 members of IF proteins with the similar structural organization (DePianto & Coulombe, 2004, Goldie, Wedig et al., 2007, Herrmann & Aebi, 2004). In the mismatched association, the complex would be dissociated into the kinked coiled-coil structures.

**Figure 7.**
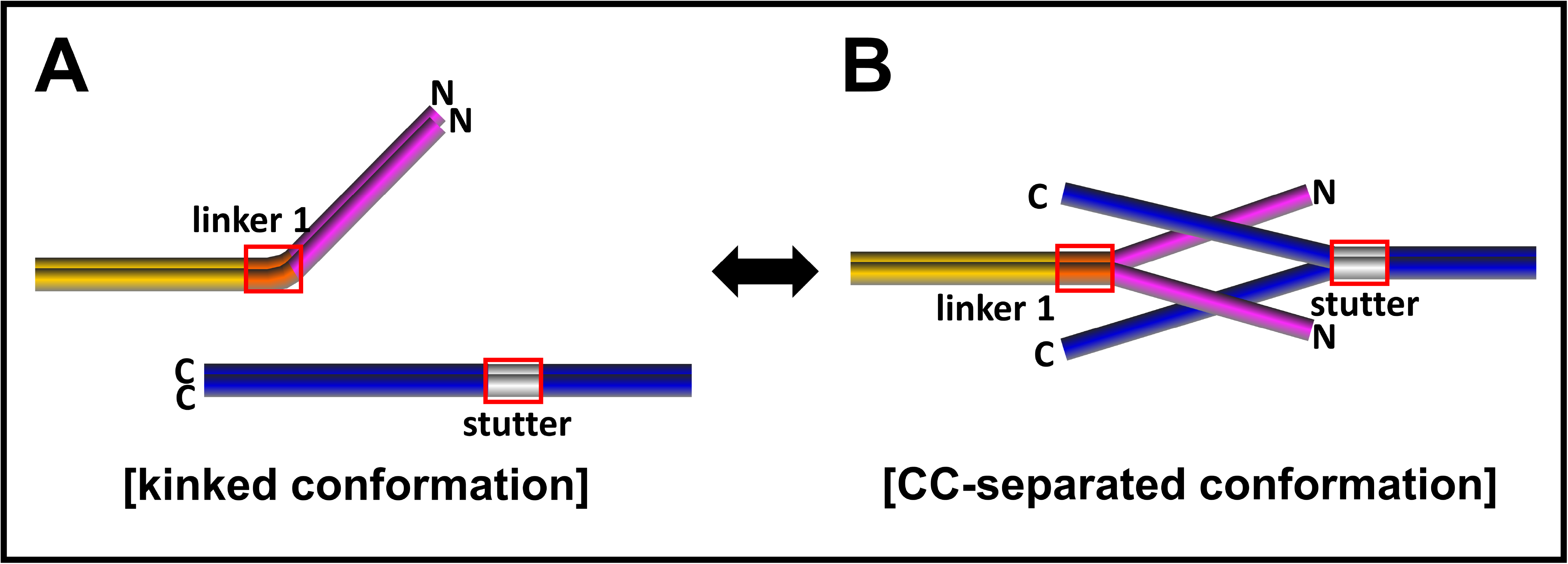
Proposed assembly model of the eA22 interaction. The kinked conformation model reflecting the lamin 300 crystal structure (Ahn et al., 2019) is shown in the left panel, and the CC-separated conformation model based on the IF assembly model (Herrmann & Strelkov, 2011, Kapinos et al., 2011) is shown in the right panel. Each subdomain of lamin A/C is coloured differently (coil 1a; magenta, linker 1; orange, coil 1b; yellow, stutter; grey, coil 2; blue). Linker 1 and stutter are shown in a red box.

Dysfunctions of coil 1a caused by genetic mutation would deteriorate dynamic remodelling or correct mesh formation in cells for the flexible and robust nuclear envelope, resulting in human diseases such as EDMD or DCM. In this study, our findings imply that the dynamic conformational changes of coil 1a and coil 2 are necessary for maintaining the normal function of nuclear lamin. Although more biochemical and genetic studies are required to demonstrate the assembly mechanism, our study provides a detailed molecular view of the structures of nuclear lamins and IF proteins in higher eukaryotes. This study will improve our understanding of the various biological processes and diseases related to IF proteins since both the weakened and augmented eA22 interactions are related to lamin-related disorders.

## METHODS AND MATERIALS

### Plasmid constructions

The genes encoding each protein were amplified DNA fragments coding for residues 1-125 (L35V, L38C, L42C, Y45C, L59R, and I63S mutants), 1-300 (L35V and I63S mutants), and 250-400 (wild-type, Y267C, K316C, L362C, L380C, L380A mutants) of lamin A/C and were inserted into the pProEx-HTa vector for overproduction of the lamin fragments (Thermo Fisher Scientific, MA, USA). The used oligonucleotide primer sequences are listed in Table EV1. The resulting plasmids encode the N-terminal His tag and the tobacco etch virus (TEV) protease cleavage site at the N-terminus of the lamin proteins. Plasmids of lamin 1-125 (wild-type), 1-300 (wild-type, Y45C, and L59R), and 250-400 were obtained, as used in previous research (Ahn et al., 2019). The plasmids were transformed into the *E. coli* BL21 (DE3) strain for overexpression. For immunofluorescence staining, the amplified DNA fragments encoding wild type and the L35V, Y45C, L59R, and I63S mutants of full-length lamin A/C were inserted into the pcDNA3.1(+) vector (Thermo Fisher Scientific, MA, USA).

### Purification of recombinant proteins

The transformed *E. coli* cells were cultured in 3 L of Terrific Broth medium containing 100 μg/ml ampicillin and 34 μg/ml chloramphenicol at 37°C to an OD_600_ of 1.0. The expression of proteins was induced using 0.5 mM IPTG at 30°C for 6 hours. After cell harvest by centrifugation, cells were resuspended in lysis buffer containing 20 mM Tris-HCl (pH 8.0), 150 mM NaCl, and 2 mM 2-mercaptoethanol. The cells were disrupted using a continuous-type French press (Constant Systems Limited, United Kingdom) at 23 kpsi pressure. The cell debris was removed after centrifugation at 19,000 g for 30 min at 4°C. The supernatant was loaded onto a cobalt-Talon affinity agarose resin (GE Healthcare). The target protein was eluted with lysis buffer supplemented with 250 mM imidazole and 0.5 mM EDTA. The lamin 250-400 protein was treated with TEV protease to cleave the His-tag. Then the target proteins were loaded onto a HiTrap Q column (GE Healthcare, USA). An increasing linear gradient of NaCl concentration was applied onto the HiTrap Q column for further purification. The fractions containing the protein were pooled and applied onto a size-exclusion chromatography column (HiLoad 26/600 Superdex 200 pg, GE Healthcare) pre-equilibrated with lysis buffer. The purified protein was concentrated and stored at −80°C until used for biochemical assays.

### Pull-down assays

A pull-down assay was conducted using His-tagged lamin proteins immobilized on the Ni-NTA resin as bait. His-tag cleaved lamin proteins as prey were incubated on the His-tagged lamin immobilized resin pre-equilibrated in a 20 mM Tris-HCl (pH 8.0) buffer containing 150 mM NaCl (or 50 mM NaCl) at room temperature for 30 min. After washing with the lysis buffer supplemented with 20 mM imidazole, the remaining resin and the fractions were analysed using SDS-PAGE. To compare the differences in binding strength of eA22 interaction with or without disulphide bond formation, experiments were performed under either the oxidized condition (0.5 mM GSSG) or the reducing condition (5 mM tris(2-carboxyethyl)phosphine; pH8.0).

### ITC

ITC experiments were conducted using an Auto-iTC200 Microcalorimeter (GE Healthcare) at the Korea Basic Science Institute (Ochang, Korea). His-tagged wild type and L35V, Y45C, L59R, or I63S mutants of lamin 300 fragments (25 μM; 0.7 mg/ml) were prepared in the sample cell (370 μL), and the TEV protease-cleaved lamin 250-400 fragment (180 μM; 3 mg/ml) was loaded into the injectable syringe. All samples were dialyzed against PBS overnight before the ITC experiments. Titration measurements of 19 injections (2 μL) with 150-sec spacing were performed at 25°C while the syringe was stirred at 750 rpm. The data were analysed using MicroCal Origin^TM^ software.

### Circular dichroism

CD spectra were collected a Chirascan plus CD spectrometer (Applied Photophysics, Surrey, UK). His-tagged lamin 1-125 fragments of wild type and L35V, Y45C, L59R, or I63S mutants in PBS (1 mg/ml) were subjected to CD experiments.

### Immunofluorescence staining

A human embryonic kidney cell line (HEK293) and human bone osteosarcoma epithelial cell line (U2OS) obtained from ATCC were maintained in liquid medium (DMEM and RPMI) containing 10% (v/v) FBS and 1% (v/v) antibiotics at 37°C. HEK293 or U2OS cells were seeded on a cover glass and transfected with the plasmid coding wild type and mutants of full-length lamin A/C using jetPEI (Polyplus Transfection). After fixing with 1% (w/v) paraformaldehyde (PFA) for 1 h at 4°C, cells were permeabilized with 0.1% (v/v) Triton X-100 for 5 min and incubated with a blocking buffer containing PBS and 5% normal goat serum (31873; Invitrogen) for 1 h. After washing with PBS twice, the cells were incubated with anti-lamin A/C (sc-376248; Santa Cruz Biotechnology) and anti-emerin (sc-15378; Santa Cruz Biotechnology) primary antibodies (1:200) in blocking buffer overnight, followed by secondary antibodies (anti-mouse Ab-FITC and anti-rabbit Ab-rhodamine; 1:400) in the blocking buffer for 7 h and mounted. The cell nuclei were stained with DAPI. The immunofluorescence signals were detected using fluorescence microscopy (Logos).

## Acknowledgments

This work was supported by grants from the National Research Foundation of Korea (2019R1A2C208513512). This research was supported by the Basic Research Program through the National Research Foundation of Korea (NRF) funded by the MSIT (2020R1A4A101932211).

---

**Table EV1.**
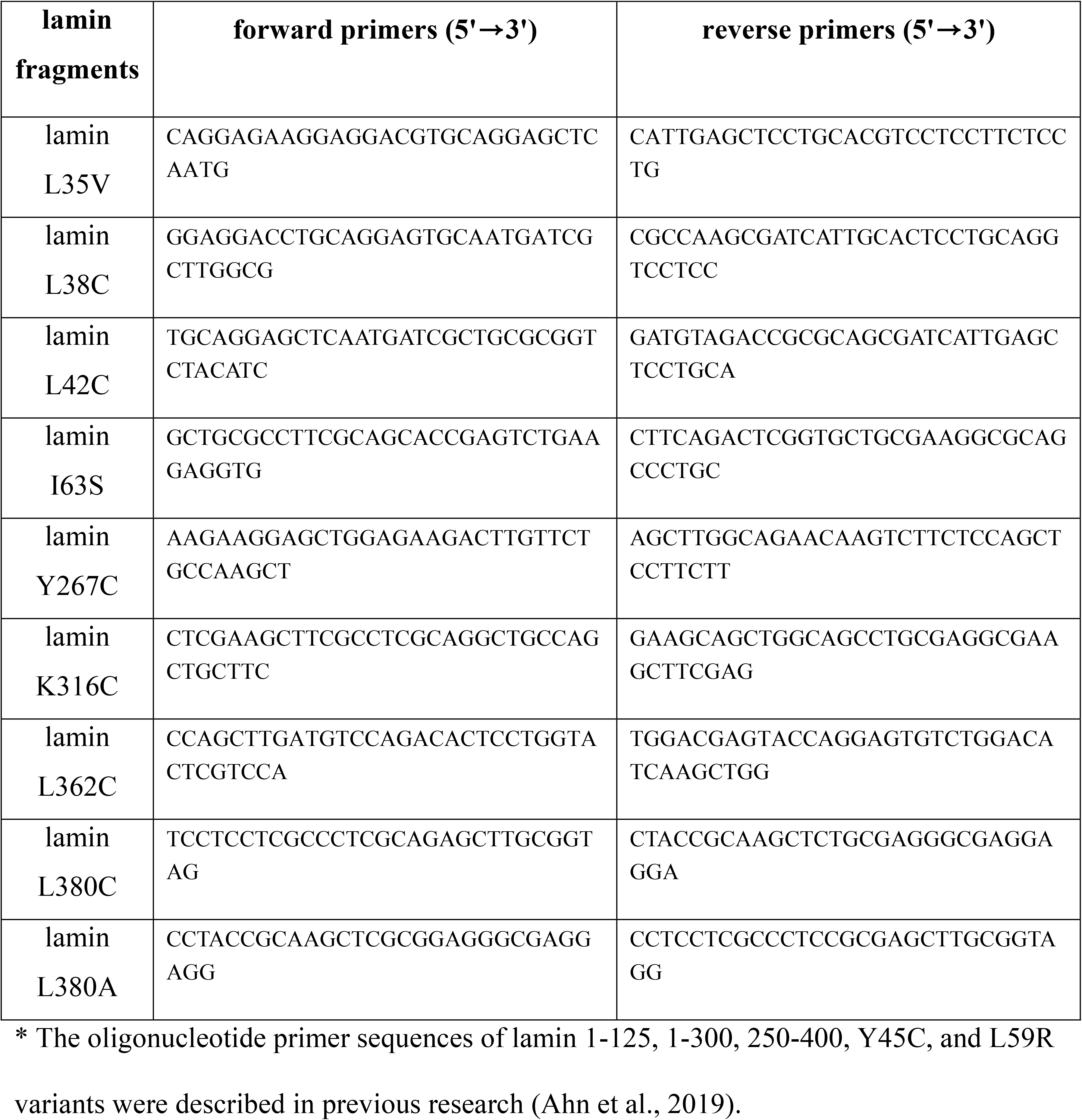
Oligonucleotide primer sequences of human lamin fragments.

**Fig EV1. Nuclear shape and distribution of lamin A/C in the human bone osteosarcoma epithelial cell line (U2OS) transfected with the plasmid encoding wild type (WT) or mutants (L35V, Y45C, L59R, and I63S) of lamin A/C**

Nuclear morphology was examined using fluorescence microscopy after transfection of wild type or the mutants (L35V, Y45C, L59R, and I63S) of lamin A/C into U2OS cells. For visualization of the nuclear membrane, the cells were stained with an anti-lamin A/C antibody for lamin A/C (green), an anti-emerin antibody for emerin (red), and DAPI for DNA (blue).

**Fig. EV2. The structural environment of L35V, Y45C, L59R, and I63S variants**

The orthogonal views of the coiled-coil structure around (A) L35V, (B) Y45C, and (C) L59R and I63S residues are presented. Each Tyr45 residue interacts with an Ile46 residue in the other protomer in the dimeric unit. Leu35, Leu59, and Ile63 residues adopt the ideal coiled-coil interaction. The predicted structural model of substituted residues (L35V, L45C, L59R, and I63S) is highlighted in red. The *a*, *d*, and *e* positions of the residues in the heptad repeat are labelled with an italic letter.

**Fig. EV3. The effects of L35V, Y45C, L59R, and I63S mutations on the binding strength between the His-tagged N-terminal fragment (residues 1-125) and the coil 2 fragment (residues 250-400) of lamin A/C**

(A) The used proteins for binding affinity were verified in SDS-PAGE.

(B) *In vitro* binding assays were conducted using the immobilized wild type or mutants of lamin 1-125 fragment on Ni-NTA resin. Lamin 250-400 fragments were incubated on empty (-) or His-lamin-bound (WT, L35V, Y45C, L59R, or I63S) Ni-NTA resins.

**Fig. EV4. The effect of the L38C and L42C mutations on binding strength of eA22 interaction**

(A) Binding affinity between the lamin 250-400 (C-terminal of coil 2) and His-lamin 1-125 fragments (WT, L38C, and L42C) was analysed by pull-down assay. To analyse the effect of the disulphide bond, oxidized (*left*), and reduced (*right*) His-lamin 1-125 fragments were used for the pull-down assay. Lamin 250-400 fragments were incubated on empty (-) or His-lamin-bound (WT, L38C, and L42C) Ni-NTA resins. The used proteins are shown in (B).

**Fig. EV5. Used proteins in the pull-down assays**

(A),(B) The used proteins in the pull-down assay shown in Fig. 6 (A; Fig 6A, B; Fig 6B).

(C) The oxidized His-lamin 250-400 proteins in Fig. 6B identified by SDS-PAGE. Protein bands for the degraded His-lamin 250-400 fragments are seen between the molecular sizes of 15 and 20 kDa in SDS-PAGE, indicated by the grey arrow.

**Fig. EV6. Straightened and CC-separated conformations of coil 1a in vimentin (PDB 3S4R (Chernyatina, Nicolet et al., 2012))**

The ribbon diagram presents the N-terminal region composed of coil 1a, linker 1, and half of coil 1b. The coil 1a region is enlarged in the bottom box. The inter-helical hydrophobic residues of coil 1b are shown as yellow sticks. The residues in the *a* and *d* positions of the heptad repeat that do not form a coiled-coil dimer are coloured magenta.

